# Training-induced improvement in working memory tasks results from switching to efficient strategies

**DOI:** 10.1101/2020.05.24.113555

**Authors:** Tamar Malinovitch, Hilla Jackoby, Merav Ahissar

## Abstract

It is debated whether training with a working memory (WM) task, particularly n-back, can improve general WM and reasoning skills. Most training studies found substantial improvement in the trained task, with little to no transfer to untrained tasks. We hypothesized that training does not increase WM capacity, but instead provides opportunities to develop an efficient task-specific strategy. We derived a strategy for the task that optimizes WM resources and taught it to participants. In two sessions, 14 participants who were taught this strategy performed as well as 14 participants who had trained for forty sessions without strategy instructions. To understand the mechanisms underlying the no-instruction group’s improvement, participants answered questionnaires during their training period. Their replies show that successful learners discovered the same strategy and improvement was associated with this discovery. We conclude that n-back training allows the discovery of strategies that enable better performance with the same WM resources.

## INTRODUCTION

Working memory (WM) is defined as the ability to simultaneously retain and manipulate information within short time periods (Baddeley, 1992b, 2003). The number of items which can be explicitly accessed and manipulated, i.e. WM capacity, is extremely limited and poses a strict bottleneck to human cognition (Cowan, 2001). Indeed, WM capacity is strongly correlated with fluid intelligence (Engle, Laughlin, Tuholski, & Conway, 1999; Süß, Oberauer, Wittmann, Wilhelm, & Schulze, 2002) and with academic achievements (Baddeley, 1992a; Bayliss, Jarrold, Baddeley, & Gunn, 2005; Hitch, Towse, & Hutton, 2001; Swanson, 2004). One of the most studied WM tasks is the n-back task (e.g. Jaeggi, Buschkuehl, Jonides, & Perrig, 2008), in which participants are presented with a sequence of serially presented stimuli and are asked to respond when a stimulus is repeated at an interval of exactly n stimuli. This task requires holding the last n items, plus the new item, in WM. When each stimulus is presented, participants must compare it to their predicted stimulus (the item presented n intervals earlier), respond if there is a match (target), and then update their WM representation to form a prediction for the next target stimulus. Since performance in this task is highly correlated with general intelligence scores, even compared with other WM tasks (Jaeggi, Buschkuehl, Perrig, & Meier, 2010), It has become a common task for training aimed at generally enhancing WM and fluid intelligence (e.g. Au et al., 2015; Redick, 2019; Schwaighofer, Fischer, & Bühner, 2015).

Training WM unequivocally yields improvement in the trained task, but the generalization of this benefit has been heatedly debated (e.g. Redick, 2019). For example, in the case of n-back, despite reports of transfer (Jaeggi, Studer-Luethi, et al., 2010), most meta-analyses and systematic reviews find no transfer to untrained tasks, or at best – minimal transfer to very similar tasks (Au et al., 2015; Melby-lervåg, 2014; Melby-Lervåg & Hulme, 2013; Melby-Lervåg, Redick, & Hulme, 2016; Redick, 2019; Soveri, Antfolk, Karlsson, Salo, & Laine, 2017). The emerging conclusion, as summarized in a recent review (Redick, 2019), is that reliable transfer of WM training occurs only when the new task is very similar (“near”) to the trained task. It seems that “far” transfer is found only in methodologically flawed studies (Melby-Lervåg et al., 2016; Redick, 2019; Sala & Gobet, 2017).

The methodological flaws appear mainly in two components of the studies (Jacoby & Ahissar, 2013, 2015; Simons et al., 2016). The first is the training protocol of the control group – far transfer has been found when a no-contact control group was included, either as the only control (e.g. Jaeggi et al., 2008) or as an additional control group, whose inclusion is crucial for attaining a significant transfer effect (e.g. Anguera et al., 2013). The no-contact group is not given monetary (or equivalent) rewards or stimulating personal attention, both of which positively impact performance. Therefore, differences in transfer may stem from the existence of a training protocol rather than from the specific training protocol of the experimental group (Foroughi, Monfort, Paczynski, McKnight, & Greenwood, 2016; Melby-Lervåg & Hulme, 2013; Shipstead, Redick, & Engle, 2012). Indeed, studies that used active control groups (trained with a similarly demanding task and a similar reward protocol) typically found either small near-only transfer (Linares, Borella, Teresa, Id, & Carretti, 2019) or no transfer at all (Jakoby, Raviv, Jaffe-dax, Lieder, & Ahissar, 2019).

The second methodological flaw is the lack of statistical correction for multiple comparisons. Typically, several tasks are assessed before and after training, and performance in most untrained tested tasks does not improve following training (Barnett & Ceci, 2002; Shipstead et al., 2012). Given that testing several tasks increases the probability of false positive(s), the target significance criterion should be raised (reviewed in Jacoby & Ahissar, 2013, 2015). However, the effect size of transfer to untrained tasks, if any, is small: ∼0.3 SD in methodologically weaker studies, ∼0.01 in methodologically sound studies (Redick et al. 2019; Melby-Lervåg, Redick, & Hulme, 2016). Since the typical size of trained groups is also small (∼ 15 per group, Chooi & Thompson, 2012; De Simoni & von Bastian, 2018; Gibson et al., 2012; Redick et al., 2013; Thompson et al., 2013), raising the target level of significance would have rendered the reported transfer non-significant (e.g. Anguera et al., 2013).

The combination of the huge effort required to conduct intensive training studies and the small (if any) generalization to untrained conditions highlights the importance of understanding the cognitive mechanisms underlying training-induced behavioral improvement. Remarkably, these mechanisms have hardly been addressed. Deciphering these processes was the aim of the current study, with a specific focus on n-back training since it is the most common trained task. We asked – what is it that participants learn which enables their substantial improvement in a challenging updating task designed to require limited-capacity online manipulations? The few recent studies that addressed this question suggest that the use of a task-specific strategy may facilitate training-induced improvement (Fellman et al., 2020; Laine, Fellman, Waris, & Nyman, 2018; Linares et al., 2019; Redick et al., 2013). Indeed, the need for a strategy that reduces WM requirements has been gradually acknowledged (Redick, 2019). But what is this strategy, and can it explain the entire learning process?

We began this study with practicing the n-back task ourselves and discussing our accumulative introspection of what facilitated our improved performance. These discussions clarified to us the strategy that we each had independently discovered. We measured whether participants who were explicitly taught this strategy could quickly reach the level of performance attained by those who go through intensive training without strategy instructions. We then deciphered, based on the self-reports of participants who had trained massively with no instructions, whether their improvement was associated with the discovery of an efficient (perhaps the same) strategy.

## METHODS

### The naïve strategy of n-updates versus the efficient strategy of 1-update

Naïve participants can typically perform well with n=1 and n=2 but find n≥3 extremely challenging. The reason the task becomes difficult with n≥3 is that participants need to update the content of n positions (slots) in WM following each item presentation. Figures 1 and 2 illustrate this naïve n-updates strategy (presented in the central column) for two types of n-back tasks – letters (Figure 1) and spatial positions (Figure 2). With this strategy, the last n items are always stored in WM in the order of their presentation. When a new stimulus is presented, it is compared to the oldest item (presented n stimuli earlier). Participants are asked to press a button if they recognize the match – stimulus repetition with an interval of n (denoted in yellow in figures 1 and 2). After each comparison, participants need to update all WM slots – all n(+1) items are “pushed” one position backward (left in Figure 1), so that the most “recent” position holds the recently presented item and the “oldest” position holds the target of the next stimulus presentation. For example, when the items are letters, n = 3, the representation in WM is D S R, and the next letter is B (Figure 1, center, line 3) – this B will be compared to the item in the slot that holds the oldest item in WM – D (center, enclosed letter), and then added to WM at the most recent slot – following R. Then the content of occupied slots in WM will be updated – shifted backwards, so that D will be dropped out, keeping the shifted 3-letter representation – S R B. Thus, the naive strategy requires an update in the content of all WM slots - a shift in the slots of all n items in memory upon each stimulus presentation.

**Figure 1:**
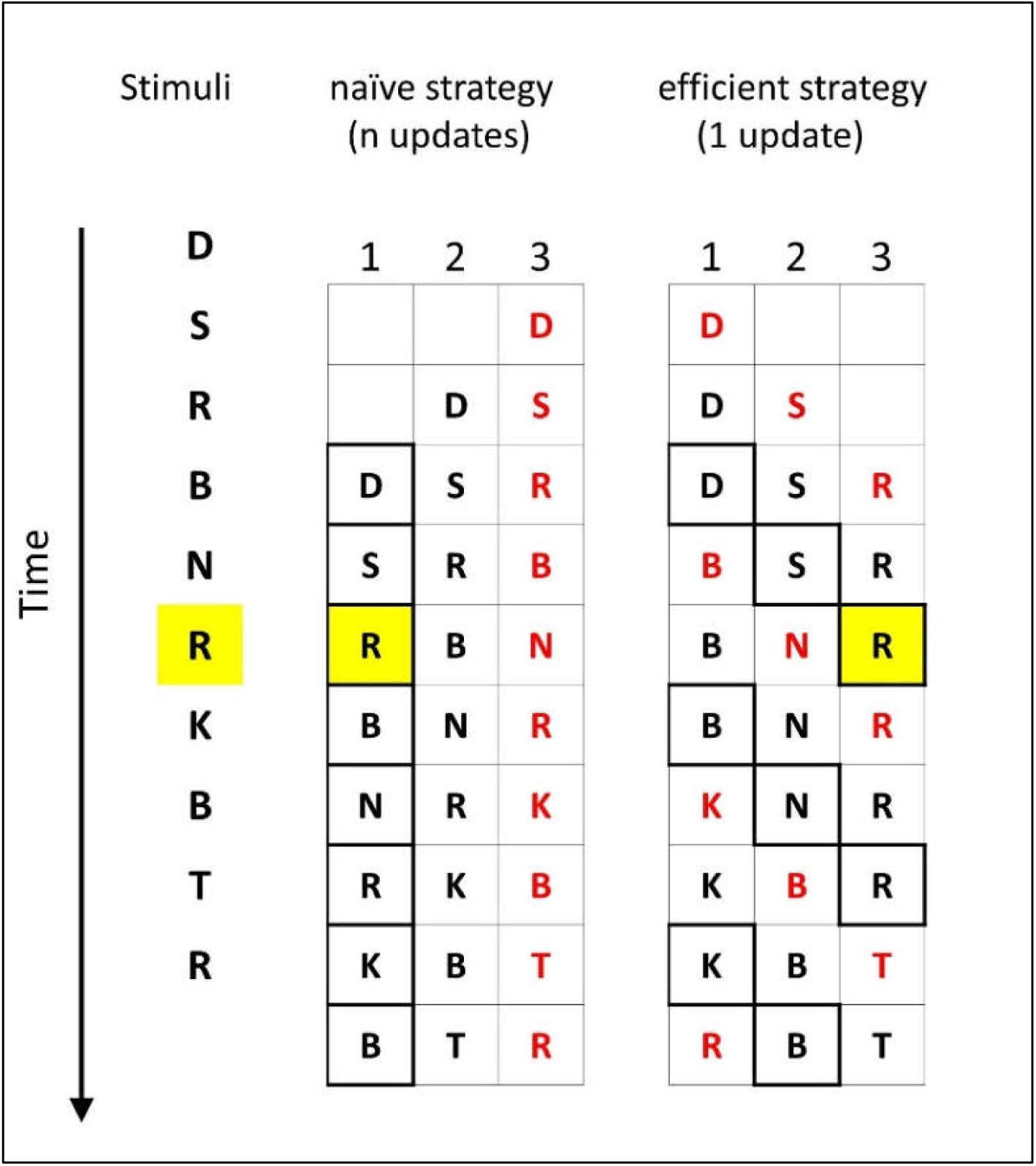
An illustration of the two strategies: naïve n-updates (center) and efficient 1-update (right) for n-back with letters, n=3. The sequence of letters is shown in the left column. Each horizontal triplet of letters represents the information stored in WM during that trial before the letter on the left is presented (after the letter above was presented). The most recent letter in each trial is denoted in red. The slot storing the content that is being compared to the incoming letter is highlighted with a bold frame. Target stimuli repeated with an interval of 3 are highlighted in yellow. The naïve strategy stores the letters in WM in the order of their presentation, and each new letter is compared to the letter that is stored in the earliest memory slot. After each comparison, all three letters that are stored in WM are shifted one slot back (the earliest letter is discarded) and the new letter is inserted into the latest WM slot. Therefore, each step requires updating the content of three slots, as with a basic stack. By contrast, in the efficient 1-update strategy, only the attended slot is updated following each new stimulus, regardless of n. The attended position is shifted every step, but there is no change in the content of unattended positions.

**Figure 2:**
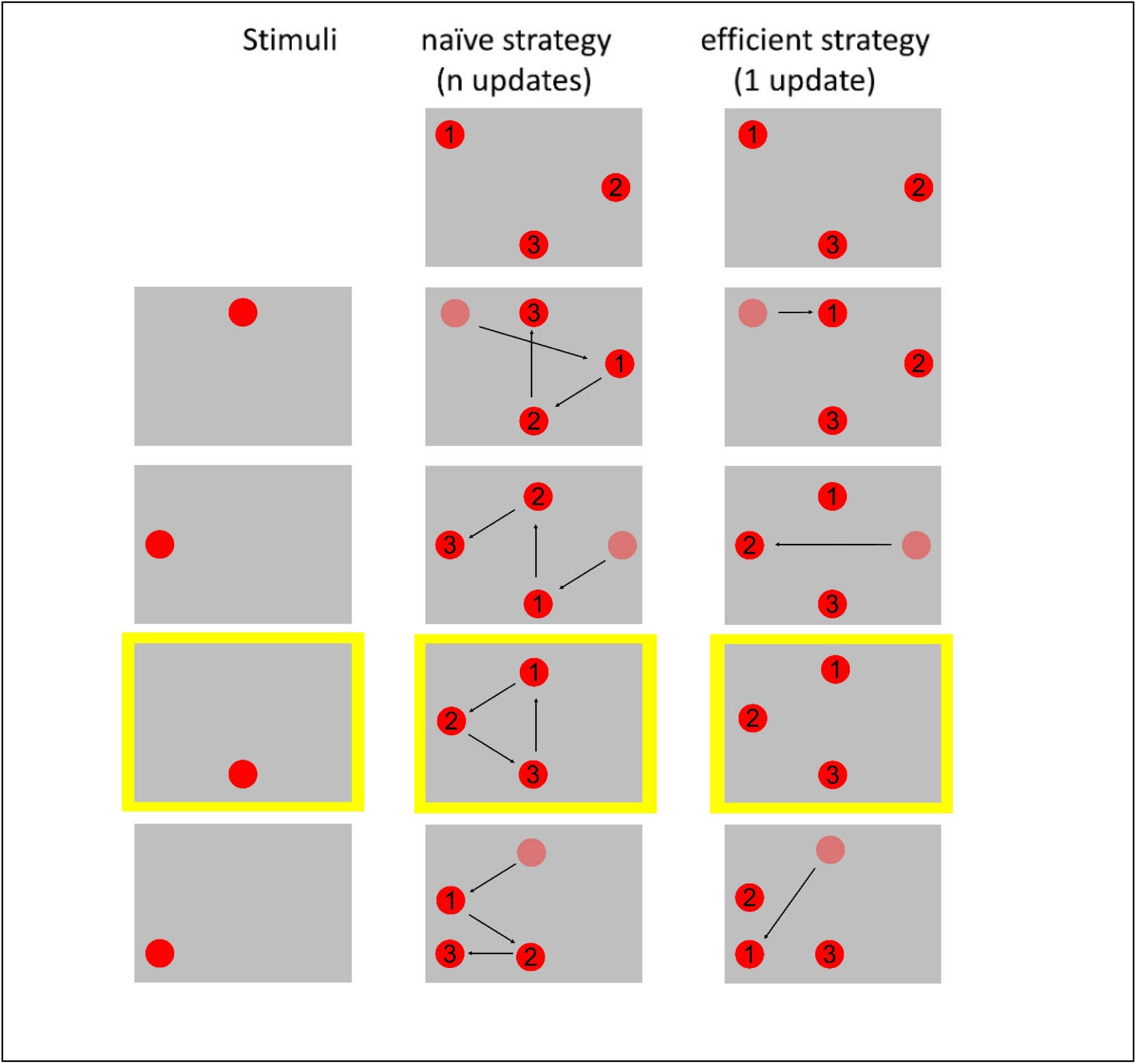
An illustration of the two strategies: naïve n-updates (center) and efficient 1-update (right) for spatial n-back, n=3. The sequence of stimuli is presented in the left column. Each triplet of circles represents the information stored in WM during a trial, when a new circle is presented. Faded red circles represent the location of the oldest stimulus in WM, soon to be deleted. The arrows represent updates of locations in WM. Repetitions with n=3 (targets) are highlighted in yellow. In the naïve strategy, locations are stored in WM in the order of their presentation. The “oldest” (presented n intervals earlier) item is compared to the newly presented item and all three slots in WM are updated, each with the content of a more recent slot – as with a basic stack. In the efficient 1-update strategy, only one WM slot is compared and updated. What changes is the attended (and updated) slot in WM. Increasing n (Figure 5) increases the tracked loop with the number of retained positions, but not the number of updates per stimulus presentation.

By contrast, the efficient 1-update strategy that we derived (Figures 1 and 2, right columns) includes no shifts in WM representation. Rather than shifting the items in WM slots, it shifts the slot to which attention is allocated in the given WM representation. The shift in the attended slot in WM does not put load on WM (Myers, Chekroud, Stokes, & Nobre, 2018). Crucially, in each trial only the item in the attended slot is updated (if it differs than the expected target). Thus, the strategy requires, at most, updating the content of one WM slot (compared with n slots in the naïve strategy), keeping track of which position is now relevant, and an attention shift, which doesn’t occupy additional WM resources. For example, for letters and n=3 (Figure 1, right), when the representation in WM is D S R, B is the new letter, and the attended position is the first (line 3), only this position is updated so that the new representation in WM will be B S R. When the next stimulus is presented (line 4), attention is shifted to the second position, and the new stimulus is compared to S, which will be updated if there is no match. Thus, if the new item is N, the updated WM sequence will be B N R. Next the third position will be attended, and then back to the first (when n=4 this loop has four positions, as illustrated in Figure 5).

**Figure 3:**
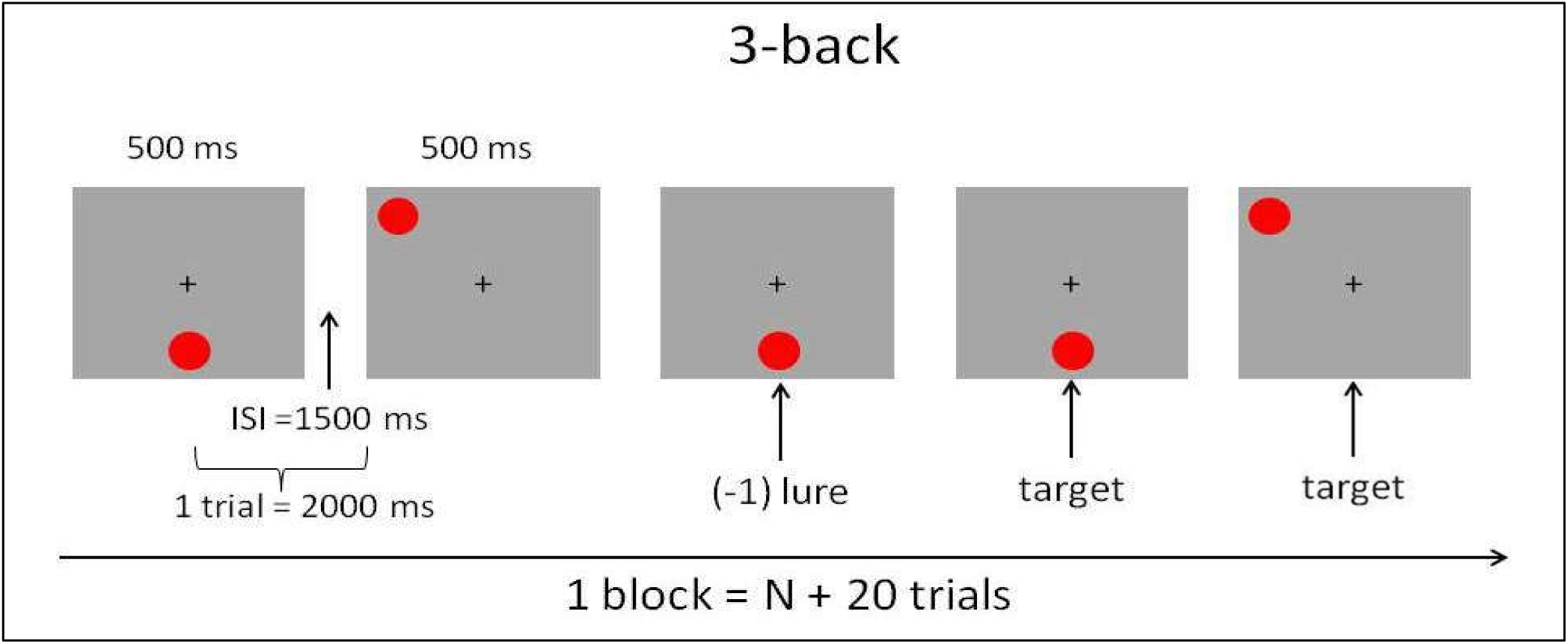
An illustration of five consecutive steps in a block of the spatial n-back task, n=3.

**Figure 4:**
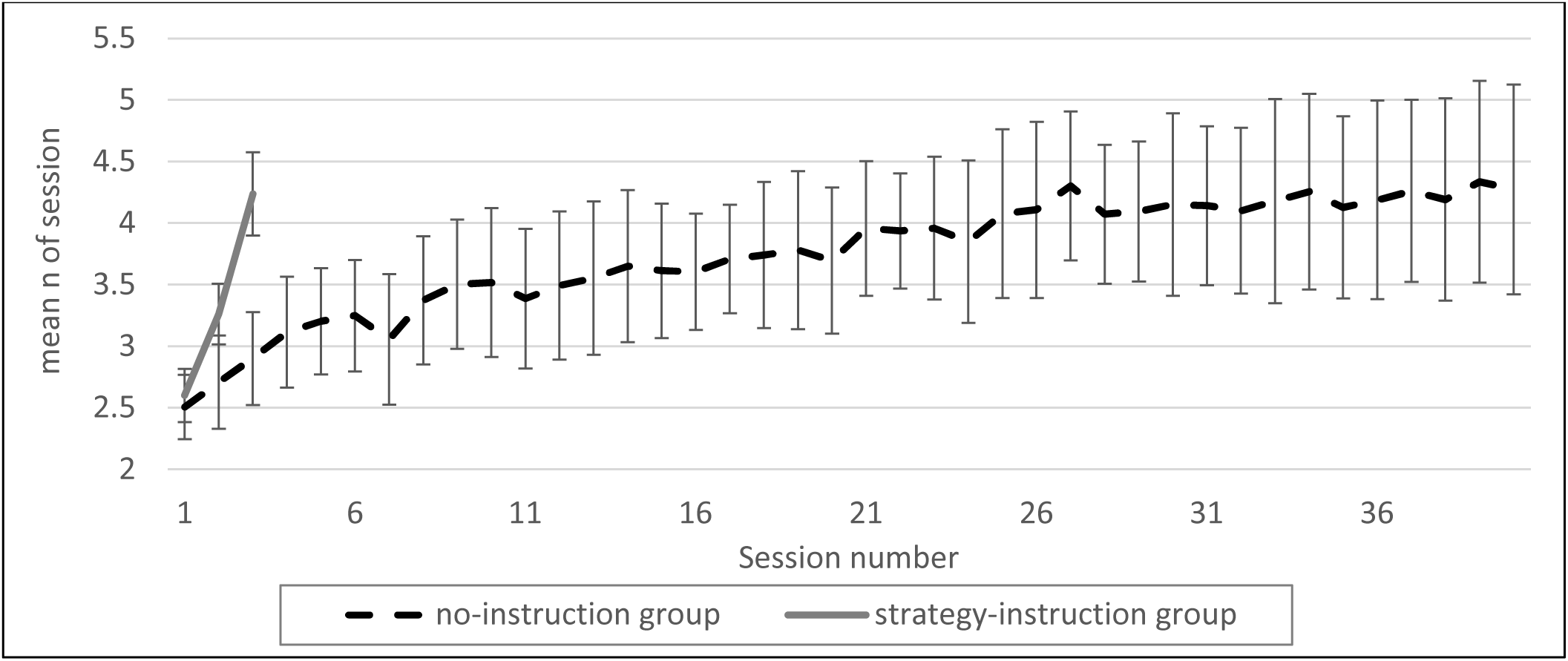
Performance as a function of session number in both groups. Mean n (∼2.5) and error bars (CI 95%) were similar for both groups during the first session. Error bars of the two groups are similar in session 1. The task is adaptive, which means that mean n increases as participants’ performance improves. Though both groups improved, the improvement rate was much faster in the strategy-instruction group. The mean n in the third session of the strategy-instruction group was similar to that attained by the no-instruction group after 25-40 sessions. The no-strategy-instruction group showed greater cross-subject variability as training progressed, revealing increased variability in learning rate. A detailed analysis of individual learning variability is presented in Figure 6.

**Figure 5:**
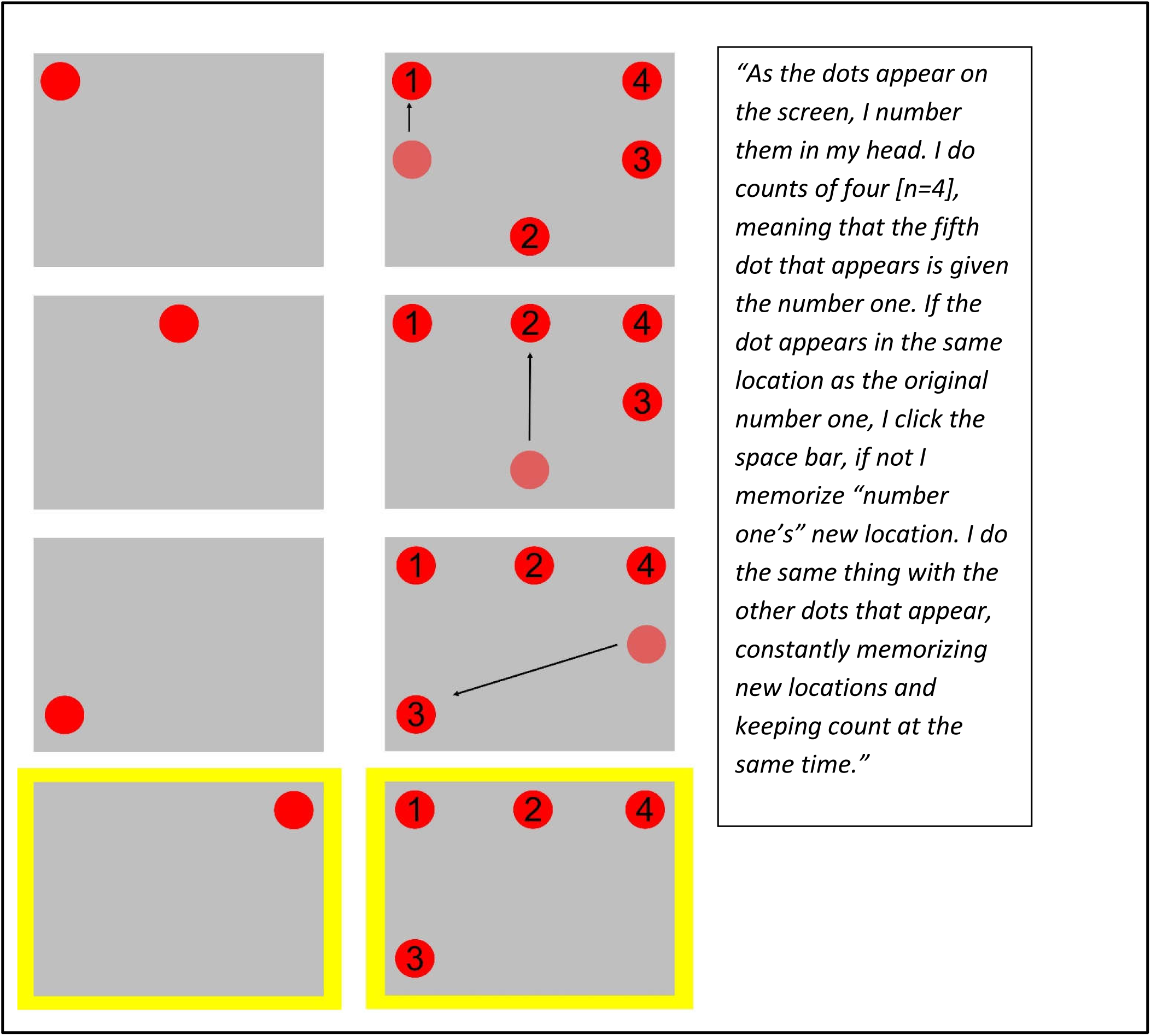
An illustration of the 1-update strategy for n=4, based on participants’ reports. Arrows represent the updated location; faded red circles represent the attended (oldest) position in WM, soon to be forgotten; a trial with a match (target) is marked in yellow.

The difference between the amount of update required by each strategy can easily be seen when examining the similarity between consecutive WM representations, shown in consecutive lines in Figure 1. One can also perceive the similarity in sound by sounding the pre- and post- updating sequences (content of consecutive steps): D S R is much more similar to B S R (efficient 1-update strategy) than to S R B (naïve strategy), since the content of only one slot is modified in the former, as opposed to three slots in the latter.

Though the above description focuses on letters, an analogous strategy can be applied to n-back with other stimuli (though this analogy may not be transparent to participants). When the task is spatial (Figure 2), spatial locations of stimuli need to be retained in WM. Thus, the same n-updates vs. 1-update strategy applies to the spatial task. In the naïve strategy, participants consistently compare the item in the first (oldest) slot, and then update the entire set of slots, by pushing them backwards and dumping the “oldest” slot from memory (as illustrated in Figure 2, left). In the efficient strategy, only one slot is updated. This slot – the attended and updated one – changes with the presentation of each stimulus, in a loop with a length of n (for n items). Here too, efficiency results from solving the task using the 1-update strategy (only one WM slot is updated in each step of Figure 2, right) and keeping track of which item needs to be attended next. As with letters, switching the strategy from updating all slots to updating only the attended slot, with the index looping over the number of items – first-second-third-first – reduces the WM resources required for attaining the same level of success.

In this study, we chose to use a spatial n-back task; in the past, we had trained a group of participants with this task, with no explicit strategy instructions, for forty sessions (Jakoby et al., 2019). Most of those participants improved significantly in this task but showed no transfer to other WM tasks. We now asked what these participants had actually learned during this training, and whether a similar degree of improvement could be gained in less time if participants were explicitly taught the efficient 1-update strategy.

### Experimental design and participants

In this paper we compared the data of two groups:

1. The strategy-instruction group (N = 14), who received three training sessions – a naïve session with no strategy instructions, and two subsequent sessions. At the beginning of each of these two sessions, they watched a detailed eight-minute video clip with strategy instructions in Hebrew (the English version of this video clip can be found here [https://youtu.be/-21tuZQNMMQ]). An experimenter asked a few questions to verify that the participants understood the strategy. Participants were then asked to perform the task according to the strategy presented in the video clip. The interval between consecutive sessions was 1-8 days. Participants were told that the aim of the study is to assess how using this specific strategy affects their performance of the task. Data for this group were collected specifically for this study.
2. The no-instruction group, who trained for forty sessions with no explicit strategy instructions (five times a week for two months). The data of this group have been previously published, in a study aimed at assessing transfer to other WM tasks, which found no transfer (Jakoby et al., 2019). Participants were told that the aim of the study is to assess how training for a task improves their performance in the trained task and in other memory-challenging tasks. Both groups answered the questionnaires detailed below, in which they expounded on the strategy they had used to perform the task.

The choice of 14 participants in the strategy-instruction group was aimed to match the number of participants who had previously formed the no-instruction group. In the context of this study, the relevant effect size is the magnitude of improvement in the trained task. Since improvement was larger than 3 SD (Jakoby et al., 2019), 14 participants per group were sufficient in both the previously studied group (no-instruction) and the newly added one (strategy-instruction group).

**Table 1:** Demographics of both groups, mean±SD; the data of the no-instruction group have been published previously (Jakoby et al., 2019).

| Group | Number (women) | Age (in years) | Years of education |
| --- | --- | --- | --- |
| Strategy-instruction | 14 (9) | 24.6 ± 2.1 | 14 ± 0.4 |
| No-instruction | 14 (11) | 23.9 ± 3.8 | 15.7 ± 2.1 |

All participants received monetary compensation or course credit for their participation (for a detailed description of the monetary compensation of the no-instruction group see Jakoby et al., 2019)). The data of one participant from the no-instruction group were excluded from the analysis because their performance on the task was an extreme outlier (z-score of over 2.5 in each session). The data of fourteen participants (out of the original fifteen) are included.

#### Spatial n-back task

Both groups were administered the same spatial n-back protocol (Jakoby et al., 2019). In this protocol, red circles are presented sequentially, one circle every two seconds (stimulus duration 500 ms; inter-stimulus interval 1,500 ms), in one of eight positions on a virtual rectangle on a computer screen. Participants respond by pressing a spacebar with their index finger whenever the location of a newly presented circle matches the location of the circle presented n steps back (target). No response is required for non-targets. Participants are notified about the relevant n at the beginning of each block. Each block comprises n+20 steps (stimuli) and includes six targets. Particularly confusing stimuli are lures: repetitions with an interval slightly different than n – a circle appears at a previous position (repetition) but with an interval of (n-1) or (n+1), as illustrated in Figure 3. Differentiating lures from targets is difficult – participants tend to press the button upon detecting a repetition, even with different intervals (Duncan, 2003). In our experiment, we included three possible levels of lure difficulty: easiest - no lures, intermediate - four lures per block (two of each type), and most difficult - eight lures per block (four of each type). We included lures because it has been previously shown that lures increase WM load and the requirements of cognitive control (e.g. Redick & Lindsey, 2013; Szmalec, Verbruggen, & Kemps, 2011). Each block’s level of difficulty was determined as follows: if the participant’s performance was 85% correct or above (calculated as hit rate minus false alarm), the difficulty level for the following block was increased by adding four more lures. After reaching a level of eight lures in a block, reaching the 85% accuracy criterion increased n by one. When performance was 65% correct or below, the number of lures was decreased from eight to four to zero, and eventually n was decreased by one (and the next block, with the smaller n, would include eight lures). Difficulty level was not modified otherwise. Each session lasted ∼30 min and consisted of 25 blocks with short breaks between them. The first two sessions began with n=2 and four lures per block for all participants. Subsequent sessions began for each participant at the difficulty level they had reached during the last block of the previous session. The same protocol was administered to both groups.

#### Questionnaires

Both groups answered questionnaires regarding the strategies they had used to perform the task. In the strategy-instruction group, participants filled out questionnaires only at the end of the third session. First, they were asked to describe their strategy in their own words, i.e. explain what they had done and evaluate the efficiency of their strategy. Then they were presented with illustrations of two strategies – the naïve n-updates strategy and the efficient 1-update strategy – and were asked to state which was closer to their own strategy (if any). This questionnaire had two goals: (1) To make sure that the participants in the strategy-instruction group had indeed used the explicitly taught strategy; (2) To see whether other methods were developed and used by the participants.

In the no-instruction group, each participant answered a questionnaire at the end of each week of training (five training sessions). The same questionnaire was administered every week. The questionnaire included wo open question regarding strategy use (“In general, could you describe your strategy for performing the training task?” and “Is it a different strategy from the one that you used in last week’s training?”). The questions were open-ended and non-specific so that no particular strategy would be implied, and no guidance would be inadvertently provided. The answers to all questionnaires were read and analyzed only after the experiment had ended, so that participants would not be affected by the experimenters’ expectations. In order to decide which strategy had been used and whether it was modified with training, we asked four independent reviewers, who were familiar with the task yet blind to participants’ performance, to evaluate based on each week’s reply of each participant whether s/he had used the efficient 1-update strategy, and if so.

## RESULTS

### Improvement was substantially faster in the strategy-instruction group

Initial performance without instructions (performance during the first session,measured by mean n per session) did not differ between groups (strategy-instruction group: mean n=2.65, SD=.37, 95% Confidence Interval (CI) 2.46 to 2.84; no-instruction group: mean n=2.43, SD=.44, 95% CI 2.2 to 2.66; p=.38 in a two-tailed, two-sample unequal variance T-test, Cohen’s d=.4).. The second session began with an instructional video clip for the strategy-instruction group, and with no specific instructions in the no-instruction group. Afterwards, both groups performed the same task, with the same adaptive protocol (see Methods section).

Mean performance in the second session significantly differed between the two groups, with the n of the strategy-instruction group (mean n=3.25, SD=.43, 95% CI 3.02 to 3.48) being significantly higher than that of the no-instruction group (mean n=2.56, SD=.58, 95% CI 2.36 to 2.82; p=.002, Cohen’s d=1.29), as shown in Figure 4.

In the third session, performance was very different between the groups (strategy-instruction: mean n=4.3, SD=.66, 95% CI 3.95 to 4.65; no-instruction: mean n=2.8, SD=.6, 95% CI 2.49 to 3.11; p<.00001, Cohen’s d=2.36). In fact, within three sessions performance of the strategy-instruction group reached the level attained by the no-instruction group only after 25-40 sessions, and did not significantly differ from the no-instruction group’s final performance during the fortieth session (strategy-instruction group third session: mean n=4.3, SD=.66, 95% CI 3.95 to 4.65; no-instruction group fortieth session: mean n=3.96, SD=1.13, 95% CI 3.37 to 4.55; p=.46, Cohen’s d=.35).

A repeated-measures two-way ANOVA for the three first sessions (2 groups X 3 sessions) showed a significant main effect of session (F(2)=62.99, p<.0001, partial ηp2=.83), indicating a general improvement as participants completed more sessions; a significant main effect of group (F(1)=17.47, p<.0001, partial ηp2=.4), indicating different performance levels in the two groups, with the strategy-instruction group showing significantly better performance overall; and crucially – a significant interaction between session and group (F(2)=26.7, p<.0001, partial ηp2=.68), indicating faster improvement in the strategy-instruction group.

Figure 4 plots mean performance (mean n per session) as a function of session number in the two groups. The initial and end points are similar, but the strategy-instruction group improved much faster. Cross-participant variability is similar for both groups in the first session (strategy-instruction SD=.37, no-instruction SD=0.44 in the no-instruction group), and it increases for both groups during training, with a greater increase observed in the no-instruction group (strategy-instruction SD=.66, no-instruction SD=1.13). This pattern results from the substantially different rates of improvement across participants, particularly when no explicit instructions are given. Large cross-participant variability was also observed in previous (no-instruction) training studies (e.g. Jaeggi, Buschkuehl, Jonides, & Shah, 2011). Previously, this variability was attributed to the extent to which general WM capacity increased. However, Figures 4 and 5 show that this cross-participant variability results from different success rates in discovering the efficient task-specific 1-update strategy.

### Improvement in the no-instruction group is associated with the discovery of the efficient strategy

Participants in the strategy-instruction group indicated in the questionnaires that they all understood and used the instructed strategy during the two post-instruction sessions.

Analyzing the self-reports of participants in the no-instruction group was complex, because the verbal reports of eight (out of fourteen) participants were too vague to determine or rule out any specific strategy. However, six (out of fourteen) participants explicitly indicated that they used the efficient 1-update strategy starting from a particular training week (see Figure 6). For example: “As the dots appear on the screen, I number them in my head. I do counts of four [n=4], meaning that the fifth dot that appears is given the number one. If the dot appears in the same location as the original number one, I click the space bar, if not I memorize “number one’s” new location. I do the same thing with the other dots that appear, constantly memorizing new locations and keeping count at the same time”. Figure 5 illustrates how this account directly maps to the implementation of the efficient 1-update strategy with n=4.

**Figure 6:**
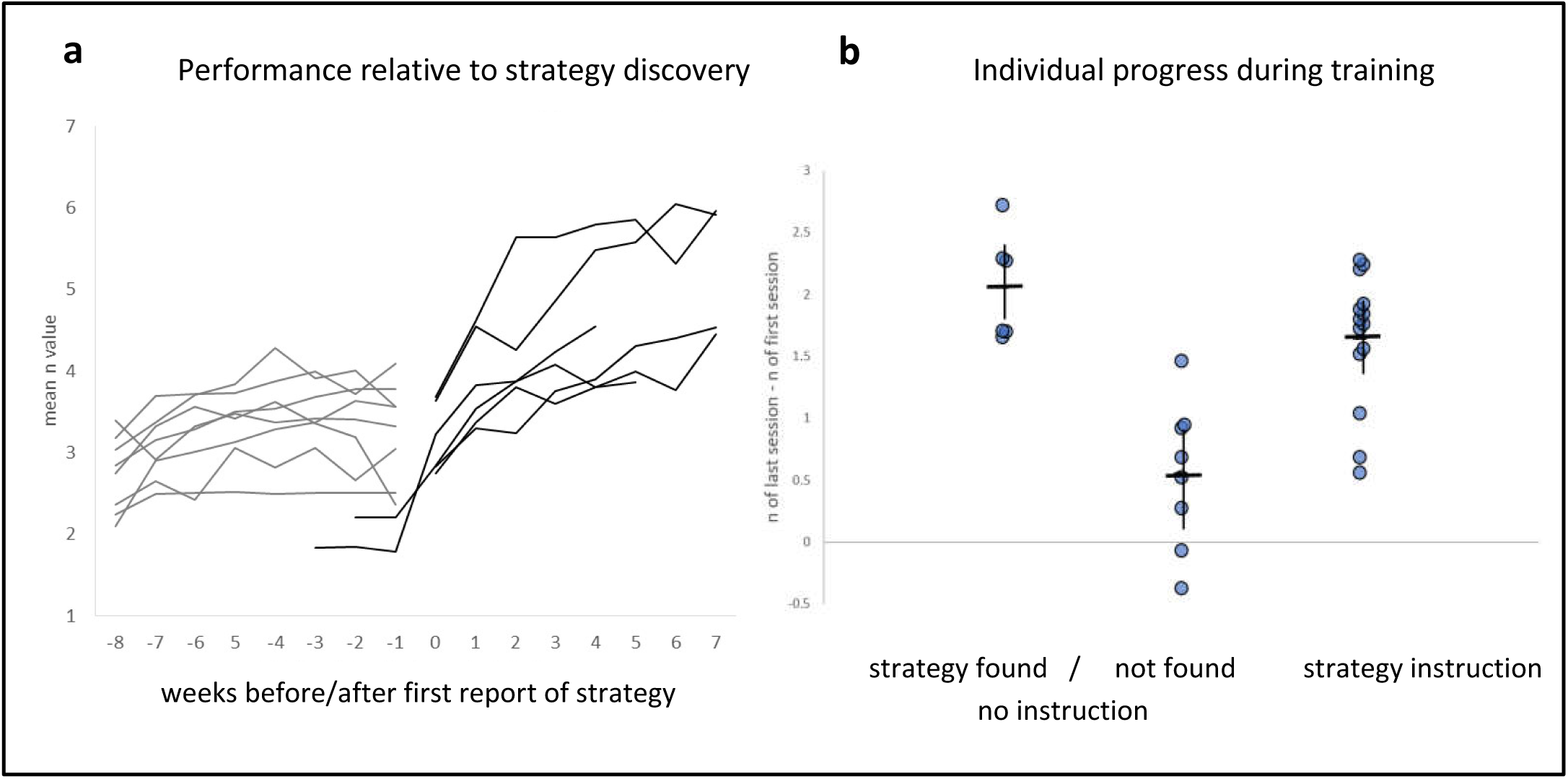
Individual learning trends. **(a)** Individual learning curves of training participants as a function of training weeks. Week 0 is the week at which they explicitly reported using the efficient 1-update strategy in their questionnaires. Grey lines represent participants whose self-reports do not indicate the discovery of an efficient strategy, and black lines represent participants who unequivocally reported the use of the strategy at some point. The mean level of n achieved by each participant is presented for each session. Explicit reports of the strategy are associated with a sharp rise in the performance curve. **(b)** Individual gains in mean-session n during training (difference between the last and first sessions) for participants in both groups. Reference lines represent group means and CI of 95%. Participants in the no-instruction group are divided into two sub-groups – those who unequivocally discovered and used the efficient strategy, and those who did not report the strategy. When examined together, the no-instruction group’s cross-participant variability in training gain is greater than that of the strategy-instruction group, in which all participant improved.

Figure 6a plots performance as a function of training week for each of the fourteen participants of the no-instruction group. Week 0 is the week at the end of which the self-report questionnaire indicated the use of the efficient 1-update strategy. As described above, the plots of six participants either started at or crossed Week 0. Four of them discovered this strategy within their first training week, i.e. within their first five sessions, and the other two discovered it later, one in the third week and the other in the fourth week. Their slopes abruptly rise following this discovery.

Figure 6b shows the individual gains in n between the first and last sessions in the strategy-instruction group and in two sub-groups of the no-instruction group – the six participants who explicitly deciphered the efficient strategy, and the eight who did not. All participants improved in the strategy-instruction group, though not to the same extent. In the no-instruction group, improvement was almost bi-modal between those who explicitly deciphered the efficient strategy and those who did not. Namely, the high cross-participant variability in the no-instruction group (Figure 4) is largely explained by the difference in improvement between those who discovered the efficient strategy and those who did not.

## DISCUSSION AND CONCLUSIONS

Our results indicate that training-induced improvement in the n-back task is due to the discovery of a task-specific strategy, rather than a general enhancement of WM capacity. This finding explains previous findings of no transfer, or only very-near transfer to other types of the n-back task (Jakoby et al., 2019; Linares et al., 2019). Performance is expected to improve in untrained tasks only when this strategy applies, and its relevance is transparent to the participants. Thus, performance in tasks with the same structure may be improved (Redick, 2019), as previously reported (Linares et al., 2019). However, performance in most other WM tasks, and even in n-back tasks for which the strategy is difficult to implement (Jakoby et al., 2019), is not expected to benefit from the discovery of this strategy.

Though to the best of our understanding the same efficient strategy was repeated across participants, we do not claim that this is the only possible strategy. Yet, theoretically, we expect all efficient strategies across WM tasks to have something in common. The number of manipulations, perhaps also the number of items accessed in WM per trial should be smaller than that of the naïve strategy. In our efficient strategy that we found this reduction is substantial. Interestingly, there is no need to reduce the total number of operations per trial, but rather only the operations that put load on WM. For example, scanning through items without modifying their slot in WM does not add to load (Myers et al., 2018). It follows that figuring out the efficient strategies that successful learners adopt will be informative regarding which operations in WM increase load versus which operations do not.

This training study specifically supports and specifies the strategy mediation theory of WM improvement, as presented by Peng & Fuchs (2017). They differentiate between the capacity theory, which claims that WM is a “mental space” that can be expanded in a muscle-like manner (Engle & Kane, 2003), and strategy mediation theory, which views WM as a finite and relatively fixed capacity, and therefore claims that WM performance is determined by the efficiency with which this capacity is used (Bailey, Dunlosky, & Kane, 2008; McNamara & Scott, 2001). According to this theory, efficient use of strategies can make more resources available for higher-level cognitive processes, which in turn can improve performance in WM tasks (McNamara & Scott, 2001; Peng & Fuchs, 2017). However, strategy mediation theory literature does not specify the nature of these putative strategies. The vague description of rehearsal or chunking (McNamara & Scott, 2001; Peng & Fuchs, 2017; Turley-Ames & Whitfield, 2003) does not capture the unique structure of WM tasks, which are designed to require online manipulations that cannot be organized into fixed chunks. The novelty of our study, from this point of view, is the ability to define a specific strategy for the well-studied n-back task and test it.

The idea that strategy plays a facilitatory role had been recently studied. For example, Felleman et al. (2020) taught participants a specific strategy for the n-back task and found that using this strategy facilitated learning. But the advantage of explicit instructions was limited, perhaps due to the internet platform they used, which did not allow them to verify whether participants understood the strategy. In our study we put a lot of effort into strategy clarification, first to ourselves and then to our participants (see video clip of instructions [https://youtu.be/-21tuZQNMMQ]). This may explain the difference in outcome. We found that strategy instruction accounted for the full magnitude of training-induced improvement. Additionally, we showed that the large individual differences in training-induced improvement within a no-strategy group delineate the participants who discovered an efficient strategy spontaneously versus those who did not.

One of the most interesting questions that our results have raised is what differentiates people who develop an efficient strategy during training, even without explicit instructions, from those who do not. Studying this question systematically requires testing whether those who developed an efficient strategy for one task also tend to develop efficient strategies for other challenging tasks, which is beyond the scope of this study. There is some evidence that individuals with larger WM pre-training are the ones who benefit more from training (Foster et al., 2017; Redick, 2019; Wiemers, Redick, & Morrison, 2019). There may be a link between initial WM capacity and the ability to quickly and efficiently adapt a strategy for a task, or there may be another cognitive trait underlying both. We do not find evidence for that in our group of participants; performance during the first session is not a good predictor of learning rate, though perhaps mean performance during the first session already includes some learning. Another suggestion in the literature is that action video game players are more likely to find efficient strategies, as their ability to learn is enhanced (Bejjanki et al., 2014, Green & Bavelier, 2014), perhaps due to enhanced attention and spatial cognition (Bediou et al., 2018). This claim had been challenged, and the findings regarding the advantages of strategic video games have been questioned (e.g. Roque & Boot, 2018; Sala, Tatlidil, & Gobet, 2018). We should note that even if action video game players demonstrate better strategies, it is still not clear whether playing action video games is the reason or the result of this enhanced strategic ability (or both).

Finally, perhaps the most important conclusion of this study is that the time has come to change the metaphors we use to describe WM training studies – like training a muscle (Jaeggi et al., 2011) or opening a bottleneck. Here we have presented an example in which the essence of WM training is not performing the same operations faster, but rather changing the set of operations used to solve the task. This type of change is likely to underly the acquisition of all expertise. When the same operations are administered to the same sequences of stimuli repeatedly, as in word reading, we replace the WM operations with chunking and schemas. But when the crux of the task is using the operations on untrained stimuli sequences (as in reading non-words), chunking cannot replace online computations. Hence, training-based improvement probably results from using a set of more efficient task-specific operations. Better understanding of these task-specific strategies may both teach us about the structure of WM and facilitate performance in tasks that heavily load on our limited WM resources.

**This work was supported by the Canadian Institutes of Health Research, the International Development Research Center, the Israeli Science Foundation, and the Azrieli Foundation (grant No. 2425/15), and by a personal grant from the Israel Science Foundation (grant No. 1650/17).**

